# ViWrap: A modular pipeline to identify, bin, classify, and predict viral-host relationships for viruses from metagenomes

**DOI:** 10.1101/2023.01.30.526317

**Authors:** Zhichao Zhou, Cody Martin, James C. Kosmopoulos, Karthik Anantharaman

**Author notes:** Correspondence (Karthik Anantharaman).

## Abstract

Viruses are increasingly being recognized as important components of human and environmental microbiomes. However, viruses in microbiomes remain difficult to study because of difficulty in culturing them and the lack of sufficient model systems. As a result, computational methods for identifying and analyzing uncultivated viral genomes from metagenomes have attracted significant attention. Such bioinformatics approaches facilitate screening of viruses from enormous sequencing datasets originating from various environments. Though many tools and databases have been developed for advancing the study of viruses from metagenomes, there is a lack of integrated tools enabling a comprehensive workflow and analyses platform encompassing all the diverse segments of virus studies. Here, we developed ViWrap, a modular pipeline written in Python. ViWrap combines the power of multiple tools into a single platform to enable various steps of virus analysis including identification, annotation, genome binning, species- and genus-level clustering, assignment of taxonomy, prediction of hosts, characterization of genome quality, comprehensive summaries, and intuitive visualization of results. Overall, ViWrap enables a standardized and reproducible pipeline for both extensive and stringent characterization of viruses from metagenomes, viromes, and microbial genomes. Our approach has flexibility in using various options for diverse applications and scenarios, and its modular structure can be easily amended with additional functions as necessary. ViWrap is designed to be easily and widely used to study viruses in human and environmental systems. ViWrap is publicly available via GitHub (https://github.com/AnantharamanLab/ViWrap). A detailed description of the software, its usage, and interpretation of results can be found on the website.

**Highlights:**

- ViWrap integrates state-of-the-art tools and databases for comprehensive characterization and study of viruses from metagenomes and genomes.
- ViWrap offers a highly flexible, modular, customizable, and easy-to-use pipeline with options for various applications and scenarios.
- ViWrap enables a standardized and reproducible pipeline for viral metagenomics, genomics, ecology, and evolution.

## INTRODUCTION

The wide application of metagenomics has deepened our understanding of the structure and function of microbiomes in mediating ecosystem processes and human health and disease. Specifically, metagenomics has offered an unprecedented window into uncultivated microbial species which are believed to account for over 99% of earth’s microbiomes [1]. The number of sequenced and publicly available metagenomes continues to increase rapidly and is enhancing our understanding of microbial communities. Though bacteria, archaea, and microeukaryotes in communities have been the primary focus of most metagenomic studies, viruses remain critically understudied. Viruses in microbial communities are typically sampled simultaneously or integrated as proviruses within microbial genomes. Since viruses are dependent on hosts for their cultivation and the vast majority of microbes remain uncultured, the study of viruses and viral communities (viromes) is being driven by metagenomics. The rapidly growing repertoire of metagenomic/viromic assemblies from various ecosystems, including natural environments, industrial man-made environments, human-microbiome related environments, etc., has provided valuable sources for mining viral diversity, studying viral roles in microbiomes, and integrating viruses into models of ecosystem function. Since 2016, scientists have greatly enriched the collection of viruses in public databases and have advanced our understanding of viruses in nature through the use of uncultivated viral genomes (UViGs) obtained from metagenomes [1]. It was discovered that viruses have significant roles in reshaping microbial host metabolism and driving global biogeochemical cycles [2, 3]. Viruses encode auxiliary metabolic genes (AMG) that augment host functions, typically for the benefit of the virus [4, 5]. These AMGs can maintain, drive, or short-circuit important metabolic steps and provide viruses with fitness advantages [5, 6]. Given the discovery of many UViGs and their AMGs, scientists have unraveled their involvement in significant ecological functions, including photosynthesis [7-9], methane oxidation [10], sulfur oxidation [11-13], ammonia oxidation [14], ammonification [15], and carbohydrate degradation [16-18], etc. In spite of these advances, our understanding of viruses continues to lag behind bacteria and archaea primarily due to the lack of available tools to study and advance viral ecology. This calls for a greater focus on the development of computational techniques facilitating virus analysis from microbiomes with a focus on metagenomic and metatranscriptomic data.

Study of viruses (involving UViGs) typically involves one of two approaches, i.e. their recovery either from bulk metagenomes or from viromes. Bulk metagenomes include all genetic materials of the microbial community, and viral fractions only account for a small portion of bulk metagenomes. Viromes, on the other hand, represent enriched and concentrated viral fractions and exclude other members of the microbial community. Many tools have been developed for the analyses of viruses based on these two approaches. VIBRANT, VirSorter2, and DeepVirFinder are three popular software for identification of viruses from bulk metagenomes and viromes. VIBRANT uses a hybrid machine learning and protein similarity approach for automated recovery and annotation of viruses [19]. VirSorter2 uses a collection of customized automatic classifiers to achieve high virus recovery performance [20]. DeepVirFinder trains viral kmer-based machine-learning classifiers to identify viruses [21].

Post virus identification, software and approaches have been developed for virus genome binning, identification of viral taxonomy, determination of genome completion estimates, and for prediction of hosts of viruses. vRhyme bins viral genomes by using both the coverage effect size and nucleotide features of viral scaffolds [22]. vConTACT2 uses whole genome gene-sharing networks for distance-based hierarchical clustering and prediction of viral taxonomy [23]. dRep enables virus clustering by dereplicating genomes based on sequence identity [24]. CheckV enables checking the quality and completeness of viral genomes [25], and iPHoP integrates all currently available virus host prediction methods and builds a machine-learning framework to obtain comprehensive host predictions for viruses [26]. Beyond these tools, multiple previously curated virus databases contain protein sequences that can be used to guide virus taxonomy classification. For example, NCBI RefSeq stores reference viral genomes [27], VOGDB provides pre-clustered viral markers of VOG HMMs (http://vogdb.org), and the IMG/VR v4 database (currently the largest virus specific genomic database) has high-quality vOTUs with taxonomy pre-assigned by stringent methods [28]. Nevertheless, these tools and databases are being increasingly used, serving as individual links within a large and complex chain of different software and approaches that are needed for comprehensive analyses of viral diversity and ecology. Given the relative infancy of the field of viromics, the knowledge of which tools to use, how to integrate methods, and to interpret results is often difficult for users with limited familiarity of viruses and bioinformatic skills. An integrated pipeline that covers the entire workflow of analyses of viruses and provides easy-to-read/parse results would significantly advance the field of virology and democratize the study of viruses from metagenomes and microbiomes.

To address this problem, we have developed ViWrap, an integrated and user-friendly modular pipeline to study viral diversity and ecology. ViWrap can identify, bin, classify, and predict viral-host relationships for viruses from metagenomes. It integrates the following advanced approaches: 1) a comprehensive screening for viruses while still keeping stringent rules; 2) a standardized and reproducible pipeline that integrates advanced tools/databases and is easy to amend for additional functionalities in the future; 3) flexible options for identifying methods, using metagenomic reads (with or without reads; short or long reads), and custom microbial genomes for various application scenarios; and 4) a one-stop workflow to generate easy-to-read/parse results with visualization and statistical summary of viruses in samples. ViWrap will significantly simplify the current computational routine to study viruses from metagenomes, speed up research in screening more viral diversity from newly generated or previously deposited metagenomes/viromes, and promote the understanding of viral community structure and function in environmental and human microbiomes.

## METHODS

ViWrap is a pipeline/wrapper to integrate several popular virus analysis software/tools to identify, bin, classify, and predict viral-host relationships from metagenomes. It takes advantage of these diverse software/tools to integrate them into a modular pipeline to obtain comprehensive information on virus genomics, ecology, and diversity in a user-friendly way. ViWrap has eight different functionalities for virus analysis including “Virus identification and annotation” (by VIBRANT, VirSorter2, and DeepVirFinder), “Virus binning” (by vRhyme), “Virus clustering” (by vConTACT2 to the genus level and dRep to the species level), “Virus taxonomy classification” (by NCBI RefSeq viral protein database, VOG HMM database, and IMG/VR v3 database [28]), “Virus information summarization”, “Result visualization”, “Virus quality characterization”, and “Virus host prediction” (by iPHoP). The intended inputs are metagenome assemblies or viromes alongside metagenomic reads. Here, we define metagenome assemblies as assemblies reconstructed from bulk metagenomes containing mixed communities of prokaryotes, eukaryotes, and viruses, and viromes as sequences from filtered/concentrated virion DNA in which viruses account for a dominant portion. Reads from metagenomes and viromes are referred to as metagenomic reads throughout the rest of the manuscript. The outputs are user-friendly tables and figures, including virus genomes and associated statistics, clustering, taxonomy, and host prediction results, annotation and abundance results, and a corresponding visualization of statistical summary (details described in Figure 1).

**Figure 1.**
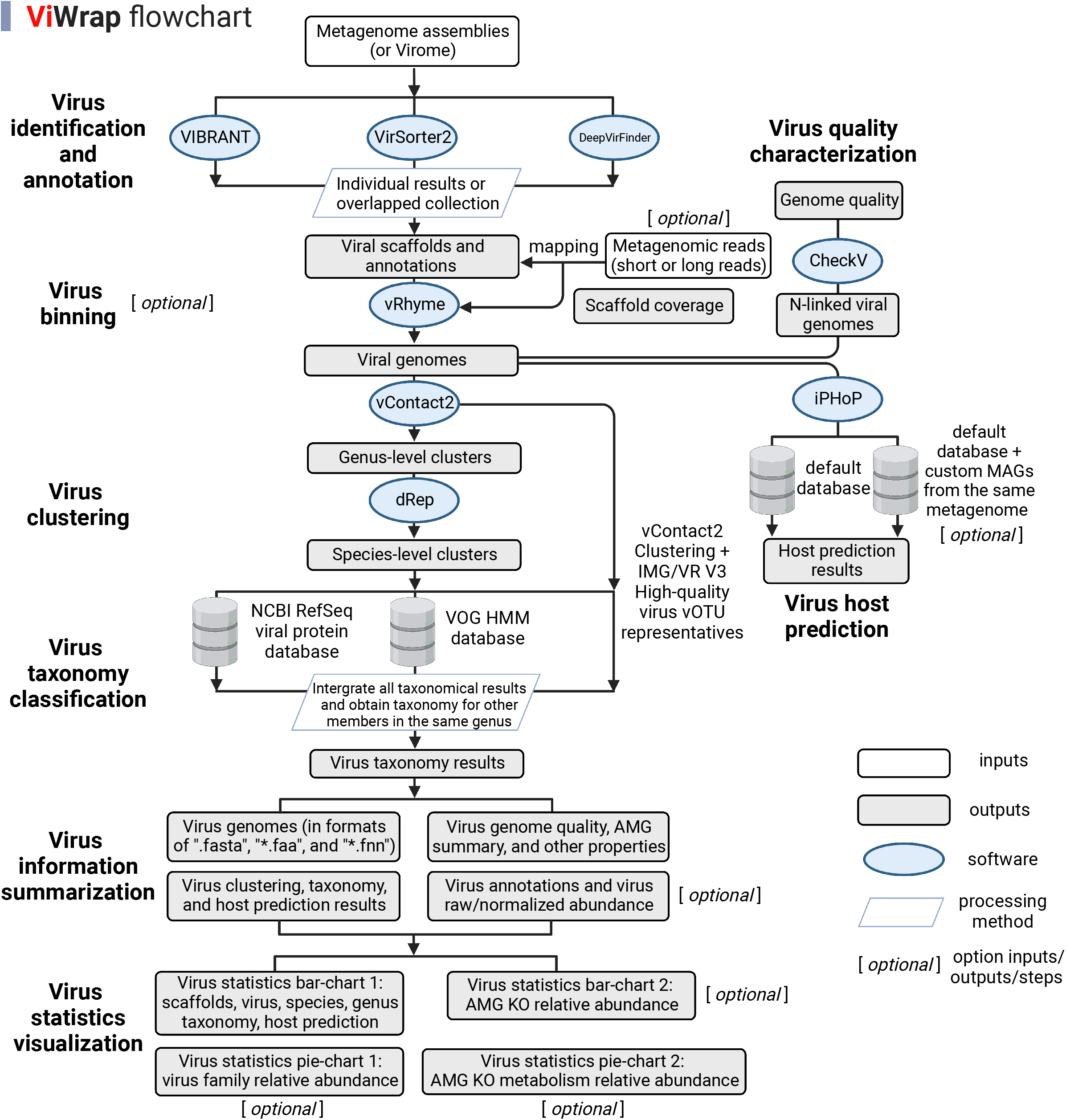
Flowchart describing the different steps and functionalities in ViWrap. Empty squares indicate inputs, filled squares indicate outputs, ovals indicate software, and parallelograms indicate the processing method that was used to get downstream results.

ViWrap can be used in conjunction with or without metagenomic reads although using reads provides advantages and enables certain analyses. Specifically, to further facilitate using metagenomes/viromes/genomes for virus mining with the corresponding metagenomic reads unavailable, we introduced a specific “run_wo_reads” python task. ViWrap is able to solely intake metagenomes/viromes or genomes without the input of metagenomic reads. When applying this task, ViWrap will avoid the steps of metagenomic mapping and virus binning, thus only reporting the results for viruses at the resolution of single scaffolds without the context of genome bins. Additionally, we implemented “set_up_env” and “download” tasks for downloading and setting up the conda environments and databases in a single step. To save on storage space required by the final result folders, we introduced a “clean” task to clean redundant information in each result directory.

ViWrap is written in Python and needs conda environments to achieve proper performance. The software is deposited in GitHub (https://github.com/AnantharamanLab/ViWrap). Details of the program’s description, installation, running methods, and explanations of inputs and outputs can be found on the GitHub page. An example ViWrap run was conducted on a metagenome dataset using the metagenomic assembly and reads of a microbial community inhabiting the deep-sea hydrothermal vent environment of Guaymas Basin in the Pacific Ocean [29]. To enable ease of use for users looking to use this as a test dataset with a shorter running time, we used a subset of the assembly (18,000 scaffolds, ∼10% of total) and two subsets of the original reads with 10% and 15% of the total reads (randomly picked) respectively as the inputs. Additionally, 98 previously reconstructed metagenome-assembled genomes (MAGs) from the same dataset were used for virus-host prediction by iPHoP based on custom host genomes.

## RESULTS

### Workflow of ViWrap

The detailed workflow of ViWrap is described in Figure 1. First, ViWrap can take metagenomic assemblies or viromes as the input source to identify viruses. Three methods were integrated to identify viral scaffolds using different algorithms, namely VIBRANT (vb), VirSorter2 (vs), and DeepVirFinder (dvf). Results of virus identification are generated using methods of a user’s choice, namely, either individual results from a single identification method (i.e., vb, vs, or dvf) or combined results by taking the intersection of results of different identification methods (i.e., vb-vs or vb-vs-dvf). These three methods have different accuracy and performance in identifying viruses. We used the “vb-vs” method as the default approach to generate a comprehensive yet stringent viral scaffold collection that meets the requirements of two popular virus identification methods (Figure 2).

**Figure 2.**
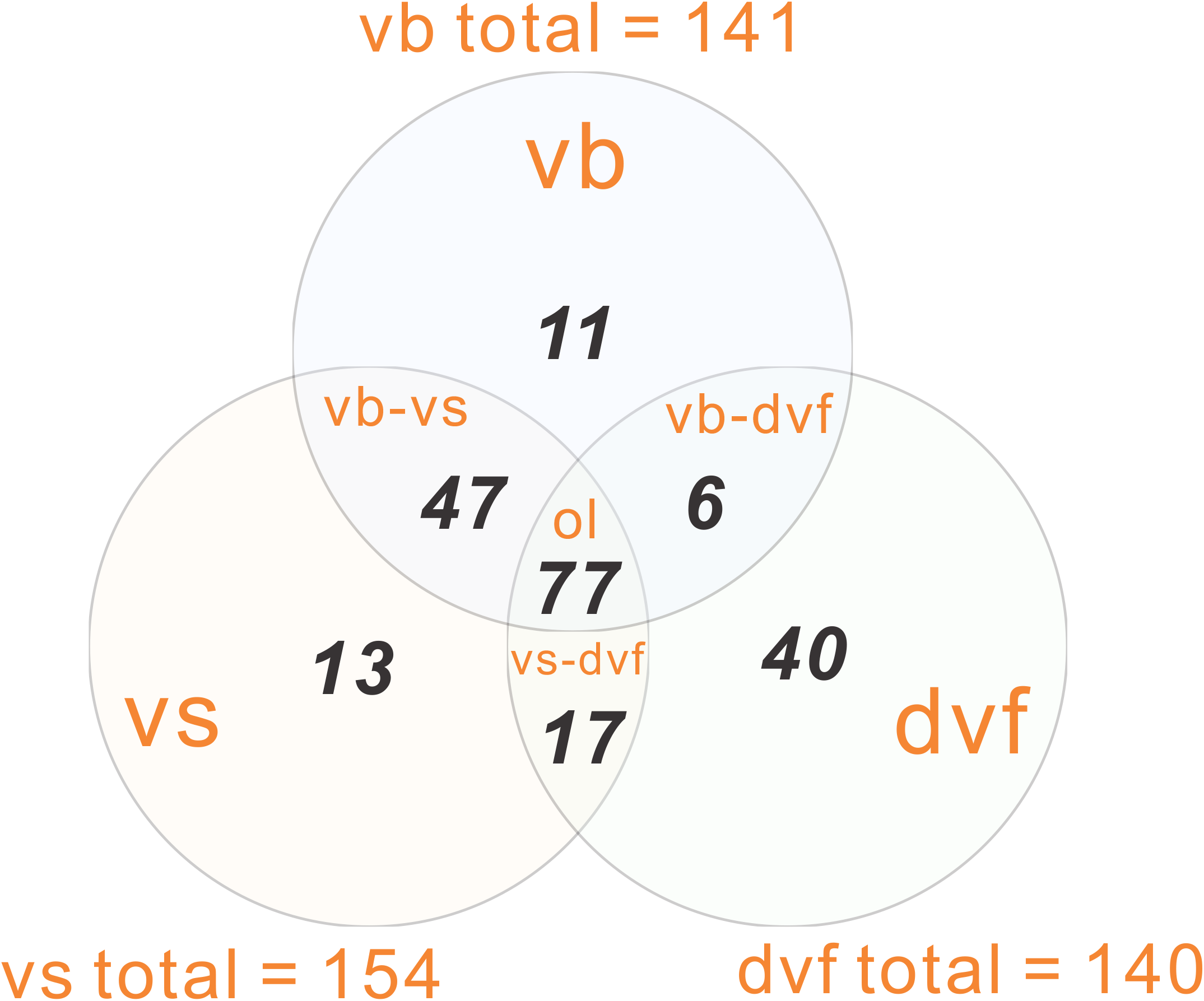
Venn diagram representing the overlapped viral scaffolds (intersection) identified by three methods. Abbreviations: “vb” – VIBRANT, “vs” – VirSorter2, “dvf” – DeepVirFinder, “vb-vs” – VIBRANT and VirSorter2, “vs-dvf” – VirSorter2 and DeepVirFinder, “vb-dvf” – VIBRANT and DeepVirFinder, “ol” – overlapped viral scaffolds by “vb”, “vs”, and “dvf”. The results of individual methods were adopted from the demonstration of example metagenome dataset of the Guaymas Basin hydrothermal vent sample.

In the second step, metagenomic reads are used to map onto the given metagenomic assemblies or viromes to get the scaffold coverage. The scaffold coverage file is used to bin viral genomes by vRhyme. To achieve stringent criteria to assign viral scaffolds into a given viral bin (viral genome), we have adopted the following requirements: 1) In vRhyme settings, the maximum protein redundancy of a viral genome was set to 5; 2) a viral scaffold that was discovered to be a “Complete” virus by CheckV is not assigned to a viral genome; 3) a bin with one or more lytic members and one integrated provirus will not be considered and will be split; 4) a bin with two or more lysogenic members (including both lysogenic scaffolds and integrated proviruses) will not be considered and will be split. Finally, CheckV is used to estimate the genome qualities of all viruses identified. Due to the fact that CheckV requires a single-scaffold virus as input, multiple fasta viral genomes were linked by multiple Ns to meet the requirement. However, because the order of linking affects ORF prediction, and some ORFs would not be called due to Prodigal’s stringency in predicting ORFs as it gets closer to the Ns junctions, these N-linked multiple fasta files are only used for estimating genome qualities by CheckV.

In the third step, genus-level clusters (viral genera) are classified by vConTACT2 (genomes within the same “VC subcluster” are regarded as from the same genus), and species-level clusters (viral species) are classified by dRep (genomes with ANI < 0.95 are regarded as from the same species).

In the fourth step, three methods are used to assign taxonomy to each virus. Two of these include protein searches using the NCBI RefSeq viral protein database and HMM marker proteins in the VOG database based on instructions described previously [28]. For the third method, we use the vOTU representatives from IMG/VR v3 high-quality vOTUs as anchors in individual genus-level clusters assigned by vConTACT2 in the previous step to assign the taxonomy information. Finally, we integrate all these three taxonomic results. When one virus has multiple taxonomic results from these three methods, the final result is provided by following the priority order of the NCBI RefSeq viral protein searching method, the VOG HMM marker searching method, and the vContact2 clustering method. To obtain the taxonomy of viruses unassigned by any of these three methods, we first enter into each genus to determine if any virus genomes have already been classified using the NCBI RefSeq viral protein searching method (only the hits from this classification method will be counted). We then expand the taxonomy to all members within the genus.

In the fifth and final step, we use iPHoP to predict hosts for viruses. Both the default iPHoP database and custom MAGs from the same metagenome can be used for host prediction. Using custom MAGs from the same metagenome can facilitate establishing direct connections between viruses and MAGs from the same community.

Finally, virus information is summarized, and statistics are visualized accordingly.

### Layout of Results

The resulting folders and files are arranged in the final output directory in the following order:

#### 00_VIBRANT_VirSorter_input_metageome_stem_name

Result of the virus identification step. This folder contains the result folders of both VIBRANT and VirSorter2 runs; additionally, a folder containing the combined results of both runs is also provided. The annotation file, “fasta” (nucleotide sequence) file, “ffn” (gene sequence) file, and “faa” (protein sequence) file are provided for viruses in the combined results.

#### 01_Mapping_result_outdir

Result of the read mapping step. Both the raw scaffold coverage result generated by CoverM (https://github.com/wwood/CoverM) and the converted coverage result used as vRhyme input are provided in the folder.

#### 02_vRhyme_outdir

Result of genome binning using vRhyme. The directory contains the folders “vRhyme_best_bins_fasta”, “vRhyme_best_bins_fasta_modified” (the best bins that were modified by stringent criteria described above), and “vRhyme_unbinned_viral_gn_fasta” (the unbinned viral scaffolds regarded as single-scaffold viruses). Additionally, it contains two tables representing the lytic/lysogenic state of viruses and genome completeness information for viruses in the “vRhyme_best_bins_fasta” folder.

#### 03_vConTACT2_outdir

Result of classification using vConTACT2. The directory contains combined protein and virus clustering results for both viruses identified from the above steps and the vOTU representatives from IMG/VR V3 high-quality vOTUs.

#### 04_Nlinked_viral_gn_dir

N-linked viral genomes used as CheckV inputs. The directory contains viral genomes with all scaffolds linked by multiple Ns. Here, only for meeting the requirement of input file format for CheckV, Single-scaffold viruses (N-linked or originally single-scaffold) are used here.

#### 05_CheckV_outdir

Result of CheckV analyses. The directory contains individual CheckV result folders for each virus and the summarized virus genome quality result with each virus as a single input.

#### 06_dRep_outdir

Result of dRep clustering. The directory contains the virus species clustering results for viruses that are assigned into the same genus.

#### 07_iPHoP_outdir

Result of host prediction using iPHoP. The directory contains the iPHoP resulting folder(s) using the default iPHoP database and custom MAGs from the same metagenome for virus identification if such custom MAGs are provided.

#### 08_ViWrap_summary_outdir

Summarized results for viruses, including “Genus_cluster_info.txt” (virus genus clusters), “Species_cluster_info.txt” (virus species clusters), “Host_prediction_to_genome_m90.csv” (host prediction result at genome level; default confidence score cutoff as 90), “Host_prediction_to_genus_m90.csv” (host prediction result at genus level; default confidence score cutoff as 90), “Sample2read_info.txt” (reads counts and bases), “Tax_classification_result.txt” (virus taxonomy result), “Virus_annotation_results.txt” (virus annotation result), “Virus_genomes_files” (containing all “fasta”, “ffn”, and “faa” files for virus genomes), “AMG_results” (containing AMG statistics and protein sequences from all virus genomes), “Virus_raw_abundance.txt” (raw virus genome abundance), “Virus_normalized_abundance.txt” (normalized virus genome abundance; normalized by 100M reads/sample), and “Virus_summary_info.txt” (summarized properties for all virus genomes, including genome size, scaffold number, protein count, AMG KOs, lytic/lysogenic state, CheckV quality, MIUViG quality, completeness, and completeness method).

#### 09_Virus_statistics_visualization

Results of visualization of Virus statistics. The directory contains two bar-charts and two pie-charts. The 1st bar-chart represents the numbers of identified viral scaffolds, viruses, viral species, viral genera, viruses with taxonomy assigned, and viruses with hosts predicted. The 2nd bar-chart represents the relative abundance of AMG KOs. The 1st pie-chart represents the relative abundance of virus families. The 2nd pie-chart represents the relative abundance of AMG KO metabolism. The raw inputs for visualization are also provided.

#### ViWrap_run.log

The log file. This file records the issued command and the time records of individual steps and the whole process.

By running the test dataset representing the Guaymas Basin deep-sea hydrothermal vent metagenome, we obtained 124 viral scaffolds that were binned into 91 viruses from the original 18,000 metagenomic scaffolds in the assembly. The total running time was ∼14 hrs using 20 threads on a Ubuntu 18.04.6 LTS (x86_64) server. For the most time-consuming parts, it took ∼2 hrs to obtain viral scaffolds from metagenomic assemblies by both VIBRANT and VirSorter2, ∼45 mins to run vConTACT2 to cluster viral genomes, ∼30 mins to conduct host prediction by iPHoP using the default database and ∼10 hrs using custom MAGs as the database (making a new database takes longer as this process is limited by the phylogenetic tree building method implemented in iPHoP).

The visualized results based on virus statistics generally represent the findings of virus numbers, taxonomy, and AMG distribution (Figure 3). From 124 viral scaffolds, 91 viral genomes (including both binned and unbinned viruses) were reconstructed (Figure 3A). Each viral genome belonged to a distinct species, and they were further classified into 81 viral genera (Figure 3A). Within the 91 viral genomes, 27 genomes had taxonomical classifications assigned, and 11 genomes had hosts predicted (Figure 3A). With regard to the taxonomy, nine families were assigned with a summed virus relative abundance of around 20.4% (Figure 3B). There were 23 AMG KOs discovered in the viral community with their corresponding relative abundance fractions assigned (Figure 3C). When classifying KOs into KEGG metabolisms, two metabolisms, carbohydrate metabolism and metabolism of cofactors and vitamins, were discovered to occupy the entire fraction (Figure 3D). The visualized results provided an intuitive and useful interpretation for general quantified features of the viral community.

**Figure 3.**
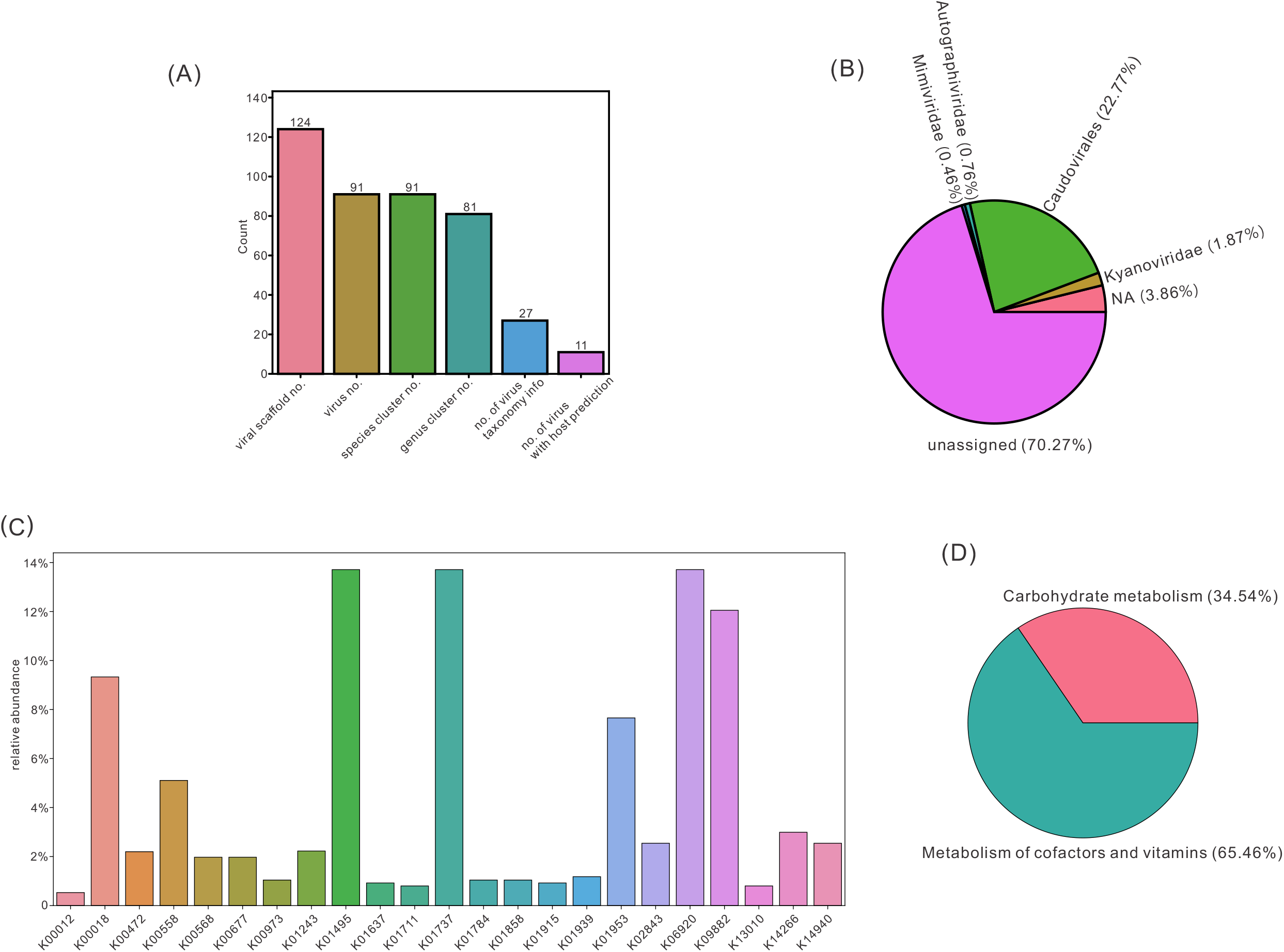
Visualizations of virus statistics. (A) Bar chart representing the numbers of identified viral scaffolds, viruses, viral species, viral genera, viruses with taxonomy assigned, and viruses with host predicted. (B) Pie chart representing the virus family relative abundance. (C) Bar chart representing the AMG KO relative abundance. (D) Pie chart representing the AMG KO metabolism relative abundance.

## DISCUSSION

ViWrap is a modular and comprehensive pipeline that integrates a full stream of virus analysis software/tools. ViWrap differs from previously developed software and tools that mainly focus on a specific “link” within the full “chain” of analyses needed for interpretation of viral diversity and ecology. Significantly, ViWrap reduces the burden on users to benchmark and choose suitable software/tools for their analyses. As the study of uncultivated viral genomes from metagenomes becomes more important [3, 30], the standardized approach of ViWrap will enable identification and analyses of viruses from metagenomes in a user-friendly manner. ViWrap integrates numerous recent mainstream and popular software/tools for virus analysis. It takes advantage of these component tools to achieve a comprehensive screening of viruses from metagenomes. The software provides flexible options for users to choose identifying methods, use metagenomic reads, and use custom MAGs from the same metagenome as an additional database for host prediction. Thus, it fits various application scenarios, i.e., unraveling viral diversity and ecology in a microbiome or environment, identifying viruses and phage in metagenomes, identifying proviruses from publicly available genomes when genomic reads are inaccessible, discovering direct connections between viruses and MAGs reconstructed from the same metagenome, etc. ViWrap also provides comprehensive virus analysis results and visualized statistics that can be easily used for further downstream analysis and interpretation of results. The summary of statistics provided by ViWrap provides a comprehensive window into the viral community and the viral ecological functions in a system.

Collectively, ViWrap is a one-stop modular pipeline and wrapper that takes metagenome/virome and/or metagenomic reads as inputs and generates easy-to-read/parse virus analysis results in a user-friendly, comprehensive, standardized (yet flexible for various application scenarios) manner. Though we demonstrate the application of ViWrap in a natural environment (hydrothermal vent environment in this study), the tools and databases implemented in ViWrap allow it to be widely used for various environments, such as man-made environmental settings (i.e., industrial environment, wastewater treatment plants), human microbiome-related environmental settings (i.e., human body, human gastrointestinal tract, oral cavity), etc. With the rapid growth of the field of virus and phage in microbiomes, larger datasets and more advanced software/tools are being developed and introduced. The modular nature of ViWrap will ensure easy integration of new tools and databases in the future. We propose that ViWrap has the potential to be widely adopted in the community and to standardize and advance the study of viruses in microbiomes.

## ACKNOWLEDGMENTS

We would like to thank members of the Anantharaman laboratory and users of ViWrap for providing useful comments and suggestions on the development of this software. This research was supported by National Institute of General Medical Sciences of the National Institutes of Health under award number R35GM143024.

## CONFLICT OF INTEREST

The authors have declared no competing interests.

## AUTHOR CONTRIBUTIONS

Zhichao Zhou and Karthik Anantharaman conceived the initial idea of ViWrap. Zhichao Zhou, Karthik Anantharaman, and Cody Martin contributed to the general function and framework of ViWrap. Zhichao Zhou and Cody Martin conducted the development and workflow of analyses. James C. Kosmopoulos contributed to the debugging process and the GitHub website. Karthik Anantharaman supervised this project. The manuscript was written by Zhichao Zhou and Karthik Anantharaman. All authors have read the final manuscript and approved it for publication.

## DATA AVAILABILITY STATEMENT

Reconstructed genomes and metagenomic reads for the example metagenome datasets from the Guaymas Basin hydrothermal vent environment are available at NCBI BioProject PRJNA522654 and SRA SRR3577362. ViWrap is publicly accessible to all researchers on GitHub (https://github.com/AnantharamanLab/ViWrap) with detailed instructions.

---

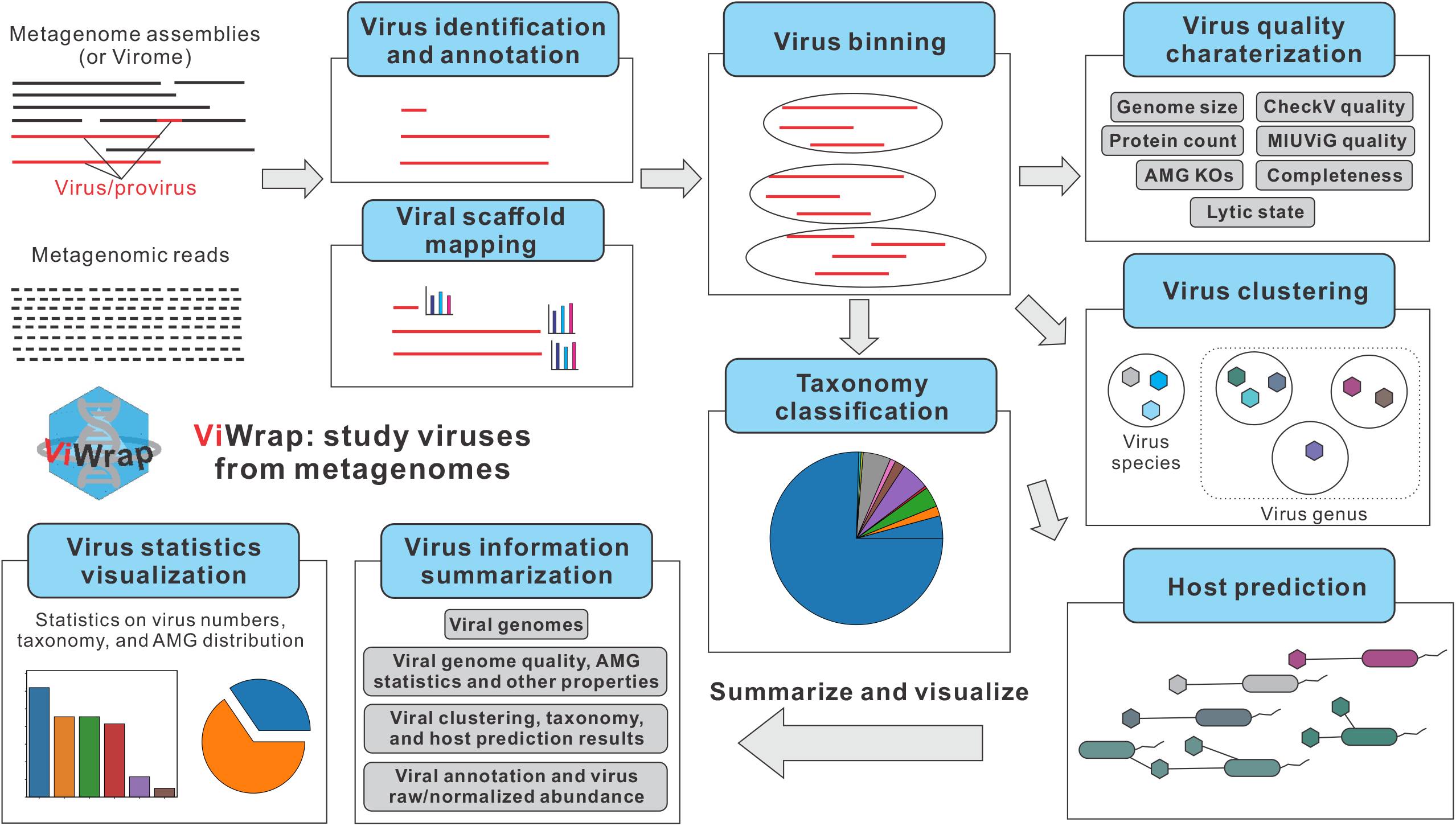

## Notes

https://github.com/AnantharamanLab/ViWrap

